# NetSurfP-2.0: improved prediction of protein structural features by integrated deep learning

**DOI:** 10.1101/311209

**Authors:** Michael Schantz Klausen, Martin Closter Jespersen, Henrik Nielsen, Kamilla Kjærgaard Jensen, Vanessa Isabell Jurtz, Casper Kaae Sønderby, Morten Otto Alexander Sommer, Ole Winther, Morten Nielsen, Bent Petersen, Paolo Marcatili

## Abstract

The ability to predict local structural features of a protein from the primary sequence is of paramount importance for unravelling its function in absence of experimental structural information. Two main factors affect the utility of potential prediction tools: their accuracy must enable extraction of reliable structural information on the proteins of interest, and their runtime must be low to keep pace with sequencing data being generated at a constantly increasing speed.

Here, we present an updated and extended version of the NetSurfP tool (http://www.cbs.dtu.dk/services/NetSurfP-2.0/), that can predict the most important local structural features with unprecedented accuracy and runtime. NetSurfP-2.0 is sequence-based and uses an architecture composed of convolutional and long short-term memory neural networks trained on solved protein structures. Using a single integrated model, NetSurfP-2.0 predicts solvent accessibility, secondary structure, structural disorder, and backbone dihedral angles for each residue of the input sequences.

We assessed the accuracy of NetSurfP-2.0 on several independent test datasets and found it to consistently produce state-of-the-art predictions for each of its output features. We observe a correlation of 80% between predictions and experimental data for solvent accessibility, and a precision of 85% on secondary structure 3-class predictions. In addition to improved accuracy, the processing time has been optimized to allow predicting more than 1,000 proteins in less than 2 hours, and complete proteomes in less than 1 day.

## INTRODUCTION

The Anfinsen experiment, showing that the structural characteristics of a protein are encoded in its primary sequence alone, is more than 50 years old (1). As a practical application of it, several methods have been developed over the last decades to predict from sequence only several protein structural features, including solvent accessibility, secondary structure, backbone geometry, and disorder (2-7). These tools have tremendously impacted biology and chemistry, and some are among the most cited works in the field. They have been extensively used to annotate novel sequences, thus facilitating their characterisation. The accuracy of said methods plays a central role here: the rate of errors in many computationally-generated annotations is a well-known and unresolved problem affecting public databases (8). 2009). Such errors often propagate through databases and sequence annotations, and the availability of high-quality predictions is hence of primary importance to limit their occurrence.

On the other hand, the amount of novel sequences has been steadily increasing over the last years (9), and not only experimental methods, but also computational predictions of structural and functional features have a hard time keeping up with it. This creates a conflict between the need for accurate predictions, and the pace at which we can generate them.

NetSurfP-1.0 (10) is a tool published in 2009 for prediction of solvent accessibility and secondary structure using a feed-forward neural network architecture. Since then, deep learning techniques have affected the application of machine learning in biology expanding the ability of prediction tools to produce more accurate results on complex datasets (11-18). Here, we present NetSurfP-2.0, a new extended version of NetSurfP, that uses a deep neural network approach to accurately predict absolute and relative solvent accessibility, secondary structure using both 3- and 8-class definitions (19), φ and ψ dihedral angles, and structural disorder (20), of any given protein from its primary sequence only. By having an integrated deep model with several outputs, NetSurfP-2.0 can not only significantly reduce the computational time, but also achieve an improved accuracy that could not be reached by having separate models for each feature. In fact, when assessed on various test sets with less than 25% sequence identity to any protein used in the training, its accuracy was consistently on par with or better than that of other state-of-the-art tools (3,4,17,21,22). In particular, we observed a significant increase in the accuracy of solvent accessibility, secondary structure, and disorder over all the other tested methods.

NetSurfP-2.0 uses different approaches to make predictions for small and large sets of sequences, thus improving its time efficiency without compromising its accuracy. It has a user-friendly interface allowing non-expert users to obtain and analyse their results via a browser, thanks to its graphical output, or to download them in several common formats for further analysis.

## MATERIALS AND METHODS

We describe briefly the dataset and method used for training NetSurfP-2.0, and the validations performed.

### Structural dataset

A structural dataset consisting of 12,185 crystal structures was obtained from the Protein Data Bank (PDB) (23), culled and selected by the PISCES server (24) with 25% sequence similarity clustering threshold and a resolution of 2.5 Å or better. To avoid overfitting, any sequence that had more than 25% identity to any sequences in the test datasets (see “Evaluation” section for details) was removed, as well as peptide chains with less than 20 residues, leaving 10,837 sequences. Finally, we randomly selected 500 sequences (validation set) left out for early stopping and parameter optimization, leaving 10,337 sequences for training.

### Structural Features

For all residues in each chain in the training dataset, we calculated its absolute and relative solvent accessibility (ASA and RSA, respectively), 3- and 8-class secondary structure classification (SS3 and SS8, respectively), and the backbone dihedral angles φ and ψ using the DSSP software (19). Finally, each residue that was present in the chain refseq sequence, but not in the solved structure, was defined as disordered. It is important to mention that disordered residues cannot be annotated with any of the other features, since no atomic coordinates are available for these residues.

### Protein sequence profiles

NetSurfP-2.0, like its predecessor, exploits sequence profiles of the target protein for its prediction. We used two different ways of generating such profiles. The first exploits the HH-suite software (25), that runs quickly on individual sequences, while the second uses the MMseqs2 software (26), that is optimized for searches on large data sets. In both cases, the profile-generation tools were run with default parameters, except MMseqs2 which used 2 iterations with the ‘--max-seqs’ parameter set to 2,000.

### Deep Network architecture

The model was implemented using the Keras library. The input layer of the model consists of the one-hot (sparse) encoded sequences (20 features) plus the full HMM profiles from HH-suite (30 features in total, comprising 20 features for the amino acid profile, 7 features for state transition probabilities, and 3 features for local alignment diversity), giving a total of 50 input features. This input is then connected to two Convolutional Neural Network (CNN) layers, consisting of 32 filters each with size 129 and 257, respectively. The CNN output is concatenated with the initial 50 input features and connected to two bidirectional long short-term memory (LSTM) layers with 1024 nodes (Figure 1, panel A).

**Figure 1.**
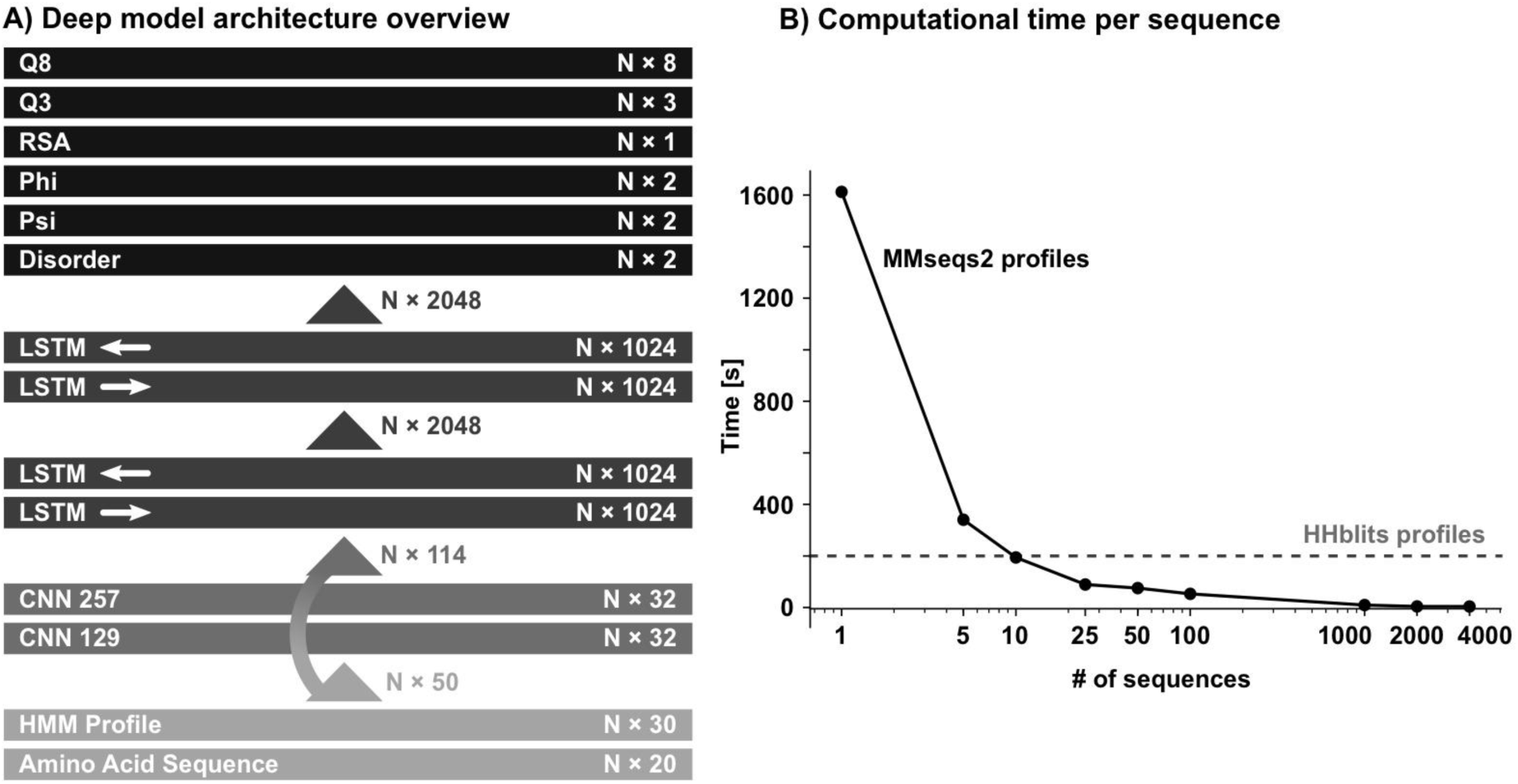
Network architecture and computation time plot. In panel the Network architecture is shown. N is the number of amino acids in the target protein sequence. Each box represents a different layer of the network, from the input (bottom) to the output (top), and the corresponding number of neurons/filters. The arrows represent the features that are passed between consecutive layers. The computation time per sequence of NetSurfP-2.0 is reported in Panel B. The x-axis represents the number of input sequences (logarithmic scale), the y-axis the average computation time in seconds per sequence. The method implementation using HH-suite profiles is reported as a grey dashed line, and the one using MMSeqs2 profiles is reported as a solid black line.

Each output (RSA, SS8, SS3, φ, ψ, and disorder) is calculated with a Fully Connected (FC) layer using the outputs from the final LSTM layer. RSA is encoded as a single output between 0 and 1. ASA output is not directly predicted, but calculated by multiplying RSA and ASAmax (27). SS8, SS3, and disorder, are encoded as 8, 3, or 2 outputs with the target encoded as a sparse vector (target is set to 1, while rest of the elements are 0). φ and ψ are each encoded as a vector of length 2, where the first element is the sine of the angle and the second element is the cosine. This encoding reduces the effect of the periodicity of the angles (28), and the predicted angle can be calculated with the arctan2 function.

### Training

The training was performed using mini-batches of size 15. The individual learning rate of each neuron was optimized using the Adam function (29). Early stopping was performed on the validation set. Since the different target values for each output have different distributions, a weighted sum of different loss functions was used: SS8, SS3 and disorder use cross entropy loss, while RSA, φ and ψ use mean squared error loss. Weights were adjusted so each loss contribution was approximately equal and then fine-tuned for maximum overall performance. When the target value for a feature of a given residue was missing, e.g. for secondary structure of disordered residues, or φ angles of N-terminal residues, the loss for that output was set to 0.

### Evaluation

The final two models one trained with the HH-suite and one with MMseqs2 profiles were tested on 3 independent datasets: the TS115 dataset (115 proteins)(Yang et al. 2016), the CB513 datasets (513 protein regions from 434 proteins) (31), and a dataset consisting of all the free-modeling targets (21 proteins) at the CASP 12 experiment (32). No protein with more than 25% sequence identity to the proteins in these datasets was present in the training. Disorder prediction was not performed on the CB513 dataset since it contains very few disordered regions.

We used different metrics to evaluate each feature: Pearson’s correlation coefficient (PCC) for solvent accessibilities, Q3 and Q8 accuracy for SS3 and SS8 respectively (17), mean absolute error in degrees for φ and ψ angles (MAE), Matthew’s correlation coefficient (MCC) and False Positive Rate (FPR) for disorder. In each dataset, the performance was calculated both as the average over all the residues in the dataset (*per residue*) and as the average of the performances per structure, the latter being defined as the average of the metric on all the residue of each given structure. The p-values between the top-scoring method and all the other methods on a given feature in a dataset were calculated using a 2-tailed paired Student’s t-test on the corresponding performances per structure.

## DISPROT dataset

We retrieved all the entries in DisProt, a database containing proteins annotated with several experimentally validated disorder types. All proteins with an available solved structure were removed to avoid overlaps with the training set. For each residue of each protein, we compare its experimental disorder annotation to the disorder prediction from NetSurfP-2.0. We classified proteins with more than 75% of their residues being disordered as completely disordered proteins.

## RESULTS

We have compared the performance of NetSurfP-2.0 to other state-of-the-art tools with similar functionality: NetSurfP-1.0 (10), Spider3 (4), SPOT-Disorder (3), RaptorX (17,22), and JPred4 (21). In order to check whether the results of the methods were significantly different, we calculated a p-value for each feature by using a pairwise Student’s t-test on the results of the two methods. Results on the independent test sets CASP12, TS115, and CB513 are presented in Table 1, and in an extended version in Supplementary table S1. They match to a very high degree with the results obtained by using a 4-fold cross validation on the training set (Supplementary table S2).

**Table 1.** Results of the method’s validation on independent test datasets. The performance of NetSurfP-2.0 (using HH-suite and MMSeqs2 profiles), NetSurfP-1.0, Spider3, SPOT-disorder, RaptorX, and JPred4, is displayed for the CASP12, TS115, and CB513 datasets. SPOT-disorder and Spider3 predictions are reported as a single row. The following performance metrics are used: Pearson Correlation Coefficient (PCC), Q3 and Q8 accuracy, Matthew’s Correlation Coefficient (MCC), False Positive Rate (FPR), and mean absolute error (MAE) in degrees. The different predicted features are reported in the column header, together with the corresponding performance metric. For each feature and each dataset, the best score is reported in bold. Scores in italics are the ones for which no significant difference with respect to the top scoring method is observed (p-value>.05). Greyed-out cells represent predictions that were not performed, either because not part of a method’s output, or because the feature was not present in the corresponding dataset.

| CASP12 | RSA<br>[PCC] | ASA<br>[PCC] | SS3<br>[Q3] | SS8<br>[Q8] | Disorder<br>[MCC] | Disorder<br>[FPR] | Phi<br>[MAE] | Psi<br>[MAE] |
| --- | --- | --- | --- | --- | --- | --- | --- | --- |
| NetSurfP-2.0 (mmseqs) | **0.726** | *0.735* | *0.820* | *0.703* | **0.660** | *0.015* | *20.3* | *31.8* |
| NetSurfP-2.0 (hhblits) | *0.725* | **0.737** | **0.824** | **0.711** | *0.604* | **0.011** | **20.0** | **31.2** |
| NetSurfP-1.0 | 0.617 | 0.641 | 0.709 |  |  |  |  |  |
| Spider3 |  | 0.688 | 0.791 |  | *0.582* | 0.026 | 21.6 | 33.2 |
| RaptorX |  |  | 0.786 | 0.661 | *0.621* | 0.045 |  |  |
| Jpred4 |  |  | 0.760 |  |  |  |  |  |
| TS115 | RSA<br>[PCC] | ASA<br>[PCC] | SS3<br>[Q3] | SS8<br>[Q8] | Disorder<br>[MCC] | Disorder<br>[FPR] | Phi<br>[MAE] | Psi<br>[MAE] |
| NetSurfP-2.0 (mmseqs) | **0.778** | **0.797** | **0.857** | **0.750** | *0.656* | **0.006** | **17.2** | **25.8** |
| NetSurfP-2.0 (hhblits) | *0.775* | *0.795* | *0.853* | *0.744* | **0.663** | *0.008* | *17.5* | *26.5* |
| NetSurfP-1.0 | 0.661 | 0.691 | 0.779 |  |  |  |  |  |
| Spider3 |  | 0.769 | 0.839 |  | *0.575* | 0.027 | 18.5 | 27.3 |
| RaptorX |  |  | 0.822 | 0.716 | *0.567* | 0.044 |  |  |
| Jpred4 |  |  | 0.767 |  |  |  |  |  |
| CB513 | RSA<br>[PCC] | ASA<br>[PCC] | SS3<br>[Q3] | SS8<br>[Q8] | Disorder<br>[MCC] | Disorder<br>[FPR] | Phi<br>[MAE] | Psi<br>[MAE] |
| NetSurfP-2.0 (mmseqs) | **0.794** | **0.807** | **0.854** | **0.723** |  |  | **20.1** | **28.0** |
| NetSurfP-2.0 (hhblits) | *0.788* | *0.803* | *0.853* | *0.720* |  |  | *20.2* | *28.6* |
| NetSurfP-1.0 | 0.700 | 0.723 | 0.788 |  |  |  |  |  |
| Spider3 |  | 0.797 | 0.845 |  |  |  | *20.4* | *28.2* |
| RaptorX |  |  | 0.827 | 0.706 |  |  |  |  |
| Jpred4 |  |  | 0.779 |  |  |  |  |  |

We give here the results for each individual feature, and a benchmark of the time performance of the tool. An example of the ASA and SS3 predictions for the human Orotate phosphoribosyltransferase (OPRTase) domain, displayed on its solved structure (PDB id 2WNS) is illustrated in Figure 2.

**Figure 2.**
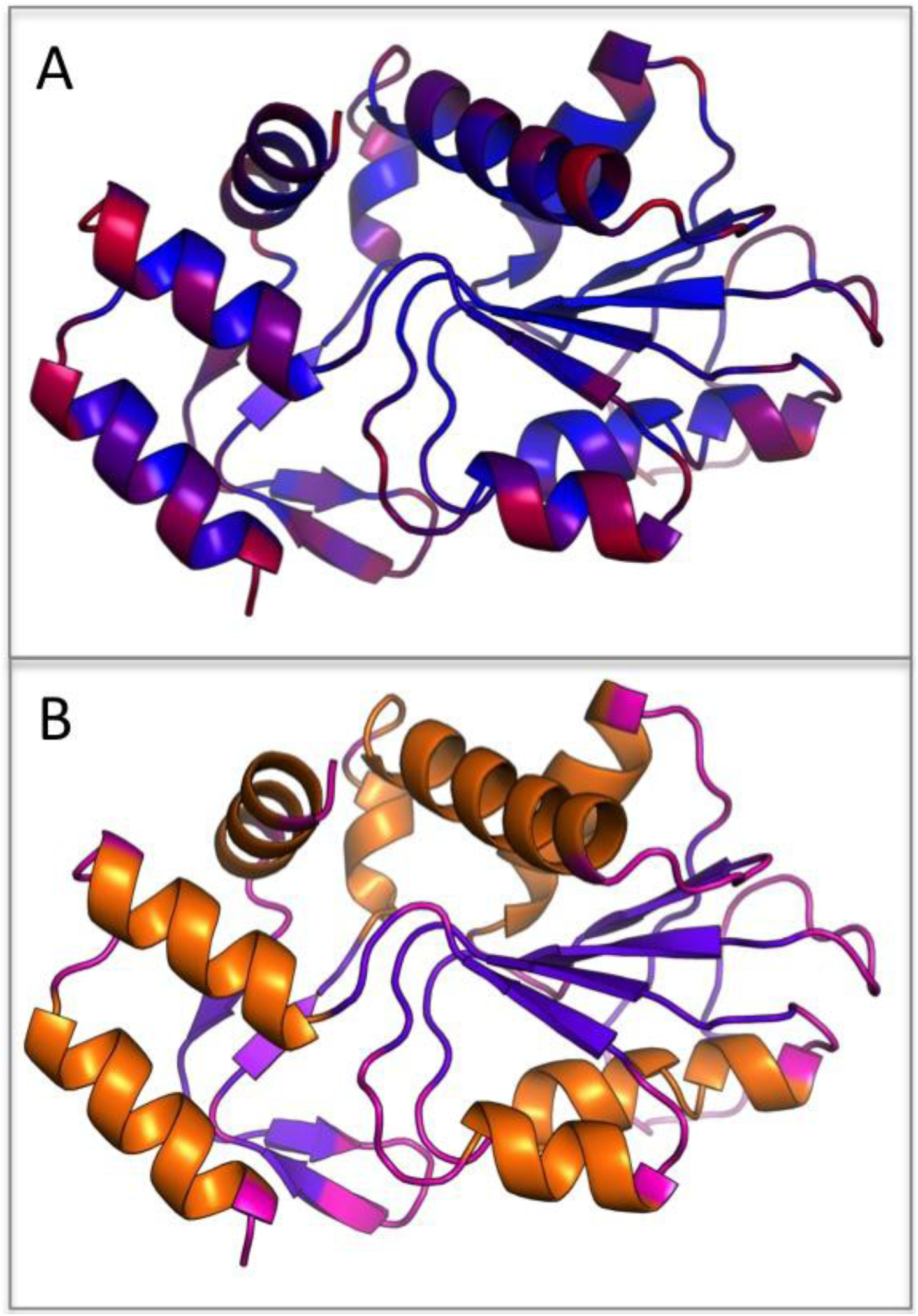
NetSurfP-2.0 predictions mapped on the OPRTase domain structure. Panel A represents the predicted ASA in a color gradient from blue (low) to red (high). Panel B represents SS3 Helix, Strand, and Coil classes in orange, purple, and pink, respectively. The actual secondary structure of the protein is displayed in the carton representation of the structure. Both color codings are consistent with the web server graphical output.

### Solvent accessibility

The main focus of the original NetSurfP-1.0 and of its updated version is to predict solvent accessibility for individual residues. Both tools predict RSA, and also calculate the corresponding ASA as described in the Methods section. We have compared the performance of the old and updated version of our tool, and compared to that of Spider3, a recent tool with a similar architecture that only predicts the ASA. The results are shown in Table 1 and demonstrate a consistently improved performance of NetSurfP-2.0, compared to the other methods, with a PCC of approximately 0.8 on the test datasets TS115 and CB513 and on the validation set. The PCC of NetSurfP-2.0 on the CASP12 dataset, though still being significantly better than all other methods, is around 0.72. It has to be noted that many of the structures in the CASP12 dataset are not obtained through X-ray crystallography, and they contain a number of disordered regions (as defined in the Methods) substantially larger than both the other external dataset and the training data. We see in general (Supplementary table S3) that all the predictions are less accurate in the few residues before and after a disordered region. We will further discuss this later.

### Secondary structure

Many tools for secondary structure prediction have been published over the last 20 years, with their reported accuracy improving over time (Yang et al. 2016). In many cases, these tools have been subject of independent validation studies (32) to get an independent assessment of their actual capabilities. We have decided to compare our tool with Spider3, RaptorX, and Jpred4, as these are among the most commonly used and accurate tools available. All the aforementioned tools perform 3-class prediction of secondary structure, while only NetSurfP-2.0 and RaptorX also provide an 8-class prediction. The results of the benchmark are given in Table 1. In all cases, NetSurfP-2.0 produce significantly more accurate predictions than all the other tools, with an average Q3 accuracy of approximately 85%, and a Q8 accuracy between 72% and 75% (Supplementary table S1, Supplementary figure S1). As for solvent accessibility prediction, the results on the CASP12 dataset are less accurate for all tested tools. A difference between the old version of the tool and it successor is that the latter is trained including disordered regions. We have also tested whether this affects the accuracy of the prediction of features other than disorder in the proximity of the disordered regions. This is actually the case for Q3 and Q8, which are significantly higher for our tool when compared to a modified version of it in which the disordered regions have been completely removed from the training sequences (Supplementary table S4). This effect is more pronounced for the datasets that have more disordered regions (CASP12 and TS115) and less so for the CB513 dataset.

Even though such residues constitute a very small portion of the total amount, these results suggest that including the disordered regions in general help our model to achieve a better internal representation of the protein sequences.

### Disorder

There is nowadays a general support that disorder plays a fundamental role in the function and dynamics of proteins, and several different types of disorder have been described and annotated. Our tool is focused on protein regions with missing atomic coordinates in their solved structures, corresponding to the REMARK-465 regions in the DisEMBL annotation (20). We have compared our prediction to both RaptorX and SPOT-disorder, a method developed by the same group that developed Spider-3. As customary in cases where the amount of negative data points greatly outnumbers the amount of positive data points (33), we have decided to use the Matthew’s correlation coefficient (MCC) to compare the tools. In all cases, NetSurfP-2.0 produces the most accurate results, with an average MCC of 0.65 (see Table 1). It has to be noted that this difference, though large, is not statistically significant.

This might be partially due to the limited number of proteins that contain disordered regions among all the datasets, and on which the MCC and the corresponding statistical tests could be calculated. On the other hand, if we extend our evaluation of the disorder prediction by including the FPR, we see that our tool produces far less false positives than all the other tools, and in this case the difference is statistically significant.

This improved performance could be due to the specific training method we used, that includes many non-disordered proteins. We will show in the following that this however does not reflect in a general underprediction tendency of the tool, as exemplified in disorder-enriched proteins.

As we mentioned above, we focus on one particular definition of disorder, while many more are available. To check if our tool can produce meaningful predictions on different types of protein disorder (e.g. intrinsically disordered proteins, functional disorder), we conducted a benchmark against the proteins in the DisProt database, a resource containing experimentally annotated disordered regions of different types. The results of this benchmark are reported in Table 2 and demonstrate that our tool performs well also on other types of disorder, and that it can also be used to identify completely disordered proteins, which were not used in its training. By comparing the recall on the TS115, the whole DisProt data set, and the subset of completely disordered protein, we see that the tool not only performs well also on proteins enriched in disordered regions, but that the low level of false positives noticed before is not linked to a general underprediction effect.

**Table 2.** Comparison of the disorder prediction on TS115, Disprot, and completely disordered proteins. The #Proteins, %disordered, and Recall column report the number of proteins, the average content of disorder, and the recall per dataset. The recall is calculated on all the disordered residues using the default 0.5 threshold.

| Dataset | #Proteins | %disordered | Recall |
| --- | --- | --- | --- |
| **TS115** | 115 | 7.4% | 51.7% |
| **DisProt Proteins** | 803 | 24.2% | 54.7% |
| **Completely Disordered Proteins** | 138 | 95.0% | 58.0% |

Another way to define disordered regions is by the so called “hot loops” (20), i.e. loop regions that present high temperature factors (B factor) for their Cα atoms. We tested if our prediction captures this definition by analysing its correlation to the B factor of backbone atoms for the TS115 dataset. To compare B factors from different structures, each B factor was normalised by the average B factor of the protein chain it belongs to. We see a Spearman correlation of 0.43 (Supplementary figure S2), confirming the ability of our model to produce a meaningful and consistent internal representation of the protein characteristics.

### Backbone dihedral angles

To complete the evaluation of our tool, we report its performance on the prediction of the backbone dihedral angles φ and ψ. We compared the results of NetSurfP-2.0 and Spider3. In this case, the two tools performed almost identically, with NetSurfP producing only marginally better predictions, with no statistically significant difference. Both tools produced more accurate prediction of the φ angle compared to ψ. This is expected, given the much broader distribution of the ψ angle in the Ramachandran plot compared to the φ angle, that is almost always confined to values between −180 and −40 degrees.

We also observed a very poor prediction accuracy for both angles in the proximity of disordered regions (Supplementary table S3), and, to a lower extent, in loop and coil regions (Table 3, Supplementary figure S3).

**Table 3:** Accuracy of phi and psi prediction according to the secondary structure for the CASP12, TS115, and CB513 datasets. In each cell we report the MAE for all residues with a specific secondary structure, either based on the 3-class (grey cells) or 8-class (white cells) definition. For the 8 class definition: G= 3-10 helix, H= □ helix, I= □ helix, B= □ bridge, E= extended strand, S= bend, T= h-bonded turn, C= coil.

|  |  | Helices |  |  | Strands |  | Coils |  |  |
| --- | --- | --- | --- | --- | --- | --- | --- | --- | --- |
|  |  | G | H | I | B | E | S | T | C |
| CASP12 | Phi | 17.35 | 8.75 | 15.72 | 29.39 | 18.43 | 34.59 | 27.65 | 28.67 |
|  |  | 9.69 |  |  | 18.89 |  | 29.91 |  |  |
|  | Psi | 30.18 | 15.51 | 16.11 | 33.73 | 20.95 | 60.73 | 33.54 | 52.54 |
|  |  | 16.93 |  |  | 21.49 |  | 50.13 |  |  |
| TS115 | Phi | 17.74 | 6.85 | 15.49 | 23.69 | 17.60 | 31.85 | 23.33 | 27.72 |
|  |  | 7.94 |  |  | 17.94 |  | 27.38 |  |  |
|  | Psi | 32.39 | 11.15 | 17.27 | 40.83 | 19.16 | 54.73 | 33.83 | 42.87 |
|  |  | 13.13 |  |  | 20.35 |  | 42.89 |  |  |
| CB513 | Phi | 19.35 | 7.88 | 16.78 | 26.14 | 17.98 | 35.67 | 27.20 | 28.66 |
|  |  | 9.24 |  |  | 18.46 |  | 29.89 |  |  |
|  | Psi | 32.66 | 10.90 | 18.14 | 40.22 | 20.18 | 56.00 | 36.35 | 41.87 |
|  |  | 13.32 |  |  | 21.36 |  | 43.64 |  |  |

### Individual vs integrated model

Thanks to the specific architecture and training strategy used, it is possible to predict all the features concurrently using a single model. Though this architecture improves the time optimization of our tool, this could potentially be sub-optimal with respect to accuracy. In order to test this, we trained single-output models for RSA, Secondary Structure (3- and 8-class), and disorder. We see (Supplementary Table S4) that the performance of the integrated model is comparable if not better than that of the individual models.

It is also interesting to notice that the hyperparameter optimisation described in Methods plays an important role in the training of the integrated model: if no relative weight is assigned to the different output, we observe a small degradation of the performance of the RSA with respect to both the integrated model and the individual ones.

### Time performance

We have shown that NetSurfP-2.0 outperforms all other methods in all the tests. This is the case for both the NetSurfP-2.0 models trained using different profile generation tools, namely HH-suite and MMseqs2. Moreover, both the HH-suite and MMseqs2 models perform similarly on all datasets tested. However, they have very different running time: the runtime on a single protein sequence for the HH-suite model is approximately 2 minutes, but it scales linearly with the number of sequences. MMseqs2, conversely, is slower for small datasets, but on large datasets it provides a speed-up of up to 50 times and the ability to parallelise on multiple processors (Figure 1, panel B). Given this, NetSurfP-2.0 is implemented to use the HH-suite model for searches of less than 100 sequences, and the MMseqs2 model otherwise, thus offering a good trade-off between computation time and resource demand, without sacrificing the method’s accuracy.

## DISCUSSION

The NetSurfP-2.0 web server provides state-of-the-art sequence-based predictions for solvent accessibility, secondary structure, disorder, and backbone geometry. By training a weight-sharing integrated model with several structural features, we improve the accuracy of disorder with respect to models trained on individual features. This improvement likely results from a more robust and informative internal state of the system, which is extremely valuable for features where only a few positives are present on average.

This integration was enabled by using improved representations of the structural data. The previous version of NetSurfP, as well as other tools, are trained only on the residues that are observed in the solved structure. In this way, the models are presented with cases that are neither physically nor biologically meaningful, such as residues divided by a disordered region, that are far apart in primary and tertiary structure but presented to the model as consecutive. In contrast, by using a recent training procedure strategy (14), we can train the model on all residues, including the disordered ones, thus increasing the accuracy of annotated features in the data and reduce the frustration during training.

The NetSurfP-2.0 framework is extremely flexible and allows to include many more structural features. We have shown that the disorder prediction of our model has a fair correlation with the residues’ B factor. Given this result, we believe that including the latter as an additional output for the system might actually improve the disorder prediction itself. Other possible features to be added are proline cis/trans conformation, metal binding sites, phosphorylation, glycosylation, and many others. Having an integrated model has an effect on the accuracy of the tool, but most importantly makes it much more time-efficient. On top of that, our software uses two different profile creation strategies in order to achieve an even better efficiency both for small sets of sequences, and for large batches of thousands of proteins. This allows the tool to annotate a single proteome in less than a day, a very important feature in present day biology.

Thanks to its accuracy, its fast computation time, and its easy and intuitive interface, we believe that NetSurfP-2.0 will become a valuable resource that will aid researchers both with and without extensive computational knowledge to analyse and understand protein structure and function.

## TOOL AVAILABILITY

NetSurfP-2.0 is available both as a web-server, and as an independent software (http://www.cbs.dtu.dk/services/NetSurfP-2.0/). The web-server version accepts up to 4,000 sequences or 4,000,000 residues per job.

## ACKNOWLEDGMENT

### FUNDING

MSK and MOAS acknowledged funding from the Novo Nordisk Foundation.

